## Supplementary Tables for "BiG-SCAPE 2.0 and BiG-SLiCE 2.0: scalable, accurate and interactive sequence clustering of metabolic gene clusters"

**Supplementary Table S1.** Extension boundaries in local alignment mode between four BGC region pairs (GBK A/B) ran with different extend strategies (Supplementary Fig. S4). Extension boundaries are represented as a slice of CDS indices (starting at 0) with an exclusive stop.

| GBK A | GBK B | Extend strategy | Ext. GBK A start | Ext. GBK A stop | Ext. GBK B start | Ext. GBK B stop |
|---|---|---|---|---|---|---|
| BGC0000253.gbk | BGC0002383.gbk | LEGACY | 9 | 17 | 2 | 10 |
| BGC0000253.gbk | BGC0002383.gbk | SIMPLE MATCH | 8 | 17 | 2 | 18 |
| BGC0000253.gbk | BGC0002383.gbk | GREEDY | 4 | 17 | 2 | 18 |
| BGC0000253.gbk | BGC0000190.gbk | LEGACY | 9 | 13 | 5 | 10 |
| BGC0000253.gbk | BGC0000190.gbk | SIMPLE MATCH | 9 | 13 | 3 | 10 |
| BGC0000253.gbk | BGC0000190.gbk | GREEDY | 3 | 22 | 0 | 29 |
| BGC0002552.gbk | BGC0002011.gbk | LEGACY | 37 | 48 | 48 | 60 |
| BGC0002552.gbk | BGC0002011.gbk | SIMPLE MATCH | 0 | 48 | 10 | 60 |
| BGC0002552.gbk | BGC0002011.gbk | GREEDY | 0 | 48 | 5 | 67 |
| BGC0001648.gbk | BGC0000086.gbk | LEGACY | 10 | 17 | 13 | 19 |
| BGC0001648.gbk | BGC0000086.gbk | SIMPLE MATCH | 3 | 22 | 4 | 21 |
| BGC0001648.gbk | BGC0000086.gbk | GREEDY | 3 | 22 | 4 | 21 |

**Supplementary Table S2.** Comparison of codebase software sustainability related metrics between BiG-SCAPE versions 1.1.9 and 2.0.0-beta.8. Lines of Code (LOC) was calculated using the pygount Python package (https://pypi.org/project/pygount/). This count is limited to python code and includes comments. Test coverage was calculated using the 'coverage' module (https://pypi.org/project/coverage/).

| **Statistic** | **BiG-SCAPE 1.1** | **BiG-SCAPE 2.0.0-beta.8** |
|---|---|---|
| LOC | 4553 | 22381 |
| Max LOC | 3333 | 1058 |
| Pylint rating | 1.74/10 | 8.75/10 |
| Count .py files | 7 | 143 |
| Tests | 0 | 400 |
| Test coverage | 0% | 77% |

**Supplementary Table S3.** Major defining characteristics of the nine benchmarking datasets with curated GCF assignments. Characteristics include dataset names and assigned codes, the number of biosynthetic regions (# BGC), the number of curated families they were grouped into (# GCF), as well as how many of these families contain only one BGC (# Singleton GCFs) and if the dataset contains manually trimmed BGC regions (Trimmed). Additionally, a count and list of the unique antiSMASH BGC classes present in the dataset is shown (# Class; Classes).

| Dataset Name | Code | # BGC | # GCF | # Singletons | # Class | Classes | Notes and other defining characteristics | Trimmed | Authors/Ref |
|---|---|---|---|---|---|---|---|---|---|
| Amycolatopsis | A | 1301 | 439 | 294 | 30 | amglyccycl, aminocoumarin, arylpolyene, bacteriocin, blactam, butyrolactone, ectoine, furan, fused, indole, ladderane, lantipeptide, lassopeptide, linaridin, melanin, nrps, nucleoside, oligosaccharide, other, otherks, phenazine, phosphonate, saccharide, siderophore, t1pks, t2pks, t3pks, terpene, thiopeptide, transatpks | BCGs derived from 28 *Amycolatopsis* strains. Large GCFs (1-43 members). Large regions, containing many hybrid classes. Contains duplicate assignments to account for assignments of neighbouring BGCs (Contains a total of 1216 unique GenBank files/regions.) | Yes | 10.1186/s12864-018-4809-4 and this publication. |
| Divergent | B | 39 | 23 | 16 | 14 | CDPS, NI-siderophore, NRP-metallophore, NRPS, NRPS-like, T1PKS, T3PKS, amglyccycl, betalactone, lanthipeptide-class-i, opine-like-metallophore, prodigiosin, siderophore, terpene | BGCs with similar gene architecture and produced compounds derived from distantly related taxa. Includes singletons from the same taxa and compound classes. See Supplementary Table S7 for further details. | Yes | This publication. Supplementary Table S7. |
| Fusarium | C | 392 | 123 | 41 | 10 | cf_fatty_acid, cf_saccharide, fatty_acid, indole, nrps, other, siderophore, t1pks, t3pks, | BCGs derived from eight *Fusarium* strains. Small curated GCFs (1-11 members). | No | 10.3389/fmicb.2018.01158 |

| | | | | | | | | | |
|---|---|---|---|---|---|---|---|---|---|
| | | | | | | terpene | | | |
| JK1 | D | 279 | 35 | 0 | 19 | NRPS, NRPS-like, PKS-like, RRE-containing, RiPP-like, T1PKS, T2PKS, T3PKS, butyrolactone, ectoine, furan, hglE-KS, indole, lanthipeptide-class-i, lanthipeptide-class-iii, lanthipeptide-class-v, melanin, siderophore, terpene | BGCs derived from nine closely related *Streptomyces* strains. Includes long-read sequencing data. Clustering was based on manual inspection of BiG-SCAPE 1 subnetworks. | No | 10.1016/j.biocontrol.2023.105411 and this publication. |
| MIBiG v1.3 | E | 281 | 90 | 12 | 34 | 2dos, LAP, NRPS, NRPS-like, PKS-like, RRE-containing, T1PKS, T2PKS, T3PKS, amglyccycl, aminocoumarin, arylpolyene, blactam, butyrolactone, cyanobactin, furan, hglE-KS, indole, ladderane, lanthipeptide-class-i, lanthipeptide-class-ii, lanthipeptide-class-iv, linaridin, microviridin, nucleoside, oligosaccharide, other, prodigiosin, resorcinol, terpene, thiopeptide, transAT-PKS, transAT-PKS-like, unknown | Grouped MIBiG 1.3 BGCs used in benchmarks of BiG-SCAPE version 1 and BiG-SLiCE version 1. | Yes | 10.1038/s41589-019-0400-9 |
| Metcalf | F | 110 | 10 | 0 | 15 | NAPAA, NRP-metallophore, NRPS, NRPS-like, PKS-like, T1PKS, T2PKS, aminopolycarboxylic-acid, betalactone, indole, lanthipeptide-class-i, | Fragmented BGCs, curated GCF assignments are based on molecular structure similarity of produced metabolites. Includes the test dataset of nine known ion signals used in the benchmarking | No | 10.1038/nchembio.1659 |

| | | | | | | | | | |
|---|---|---|---|---|---|---|---|---|---|
| | | | | | | lassopeptide, nucleoside, terpene, transAT-PKS | of BiG-SCAPE version 1 parameters. | | |
| Planomonospora | G | 11 | 3 | 0 | 5 | LAP, NRPS, lanthipeptide, thiopeptide, unknown | Very small dataset. Validated using metabolomics. | No | 10.1021/acs.jnatprod.0c00807 |
| Reclassification | H | 90 | 10 | 1 | 4 | NRPS, NRPS-like, T3PKS, phosphonate | Highly similar (gene architecture wise) glycopeptide NRPS BGCs. Curated GCF assignments group these into subtypes. A number of these BGCs also include neighbouring T3PKS/phosphonate protoclusters. | No | 10.1101/2023.02.10.526856 and this publication. |
| Rhodococcus | I | 408 | 34 | 1 | 11 | butyrolactone, cf_fatty_acid, cf_putative, cf_saccharide, ectoine, fatty_acid, nrps, other, saccharide, t1pks, terpene | Large GCFs (1-53 members) curated based on manual MultiGeneBlast analyses per family. | No | 10.1186/s12864-017-3966-1 |

**Supplementary Table S4.** Curated GCF assignments for benchmarking dataset B (Divergent). Seven GCFs contain BGCs that produce the same or highly similar compounds in relatively distant taxa. An additional 16 unrelated singleton GCFs were added as negative signals spanning across these taxa and compound classes.

| BGC | GCF number | taxonomy | Notes | Accession | Source |
|---|---|---|---|---|---|
| BGC0000453.region001 | 1 | *Streptomyces tsusimaensis* | Produces valinomycin | | MIBiG |
| BGC0001846.region001 | 1 | *Streptomyces sp. CBMAI 2042* | Produces valinomycin | | MIBiG |
| BGC0001341.region001 | 1 | *Rothia nasimurium* | Produces valinomycin | | MIBiG |
| BGC0001468.region001 | 2 | *Streptomyces cinnamoneus* | Produces bicyclomycin | | MIBiG |
| NC_023149.1.region016 | 2 | *Pseudomonas aeruginosa SCV20265* | Produces bicyclomycin | GCF_000510305.1 | 10.1128/AEM.02828-17 |
| BGC0000825.region001 | 3 | *Streptomyces sp. TP-A0274* | Produces staurosporine | | MIBiG |
| BGC0000826.region001 | 3 | *Streptomyces clavuligerus ATCC 27064* | Produces staurosporine | | MIBiG |
| BGC0000827.region001 | 3 | *Salinispora arenicola CNS-205* | Produces staurosporine | | MIBiG |
| BGC0001453.region001 | 4 | *Streptomyces argillaceus* | Produces desferrioxamine | | MIBiG |
| NC_019936.1.region003 | 4 | *Pseudomonas stutzeri RCH2* | Produces desferrioxamine | NC_019936.1 | 10.1093/femsec/fix086 |

| | | | | | |
|---|---|---|---|---|---|
| NC_017390.1.region004 | 4 | *Erwinia pyrifoliae DSM 12163* | Produces desferrioxamine | NC_017390.1 | 10.1093/femsec/fix086 |
| BGC0001572.region001 | 4 | *Pantoea agglomerans* | Produces desferrioxamine | | MIBiG |
| NZ_JFGV01000084.1.region001 | 4 | *Photorhabdus luminescens BA1* | Produces desferrioxamine | GCF_000612035.1 | 10.1093/femsec/fix086 |
| NC_014228.1.region003 | 4 | *Xenorhabdus nematophila ATCC 19061* | Produces desferrioxamine | NC_014228.1 | 10.1093/femsec/fix086 |
| BGC0000259.region001 | 5 | *Serratia marcescens* | Produces prodigiosin | | MIBiG |
| BGC0000258.region001 | 5 | *Serratia sp.* | Produces prodigiosin | | MIBiG |
| NZ_CP045429.1.region009 | 5 | *Pseudoalteromonas rubra S4059* | Produces prodigiosin | NZ_CP045429.1 | 10.1128/Spectrum.01171-21 |
| BGC0000260.region001 | 5 | *Hahella chejuensis KCTC 2396* | Produces prodigiosin | | MIBiG |
| BGC0000137.region001 | 6 | *Salinispora arenicola CNS-205* | Produces rifamycin | | MIBiG |
| BGC0000136.region001 | 6 | *Amycolatopsis mediterranei S699* | Produces rifamycin | | MIBiG |
| BGC0002412.region001 | 7 | *Photobacterium galatheae* | Produces holomycin | | MIBiG |
| BGC0002091.region001 | 7 | *Yersinia ruckeri ATCC 29473* | Produces holomycin | | MIBiG |

| | | | | | |
|---|---|---|---|---|---|
| BGC0000373.region001 | 7 | *Streptomyces clavuligerus ATCC 27064* | Produces holomycin | | MIBiG |
| NC_023149.1.region006 | 8 | *Pseudomonas aeruginosa SCV20265* | NRPS Singleton | GCF_000510305.1 | NCBI |
| NC_014228.1.region010 | 9 | *Xenorhabdus nematophila ATCC 19061* | NRPS Singleton | NC_014228.1 | NCBI |
| NC_014228.1.region001 | 10 | *Xenorhabdus nematophila ATCC 19061* | NRPS Singleton | NC_014228.1 | NCBI |
| NC_017390.1.region002 | 11 | *Erwinia pyrifoliae DSM 12163* | NRPS Singleton | NC_017390.1 | NCBI |
| NZ_RCOL01000001.1.region030 | 12 | *Streptomyces sp. CBMAI 2042* | NRPS Singleton | NZ_RCOL01000001.1 | NCBI |
| NC_023149.1.region009 | 13 | *Pseudomonas aeruginosa SCV20265* | NRPS Singleton | GCF_000510305.1 | NCBI |
| NC_023149.1.region015 | 14 | *Pseudomonas aeruginosa SCV20265* | Opine-like metallophore singleton | GCF_000510305.1 | NCBI |
| NZ_AEDL01000001.1.region002 | 15 | *Pantoea sp. aB* | Terpene singleton | GCF_000179655.1 | NCBI |
| NZ_RCOL01000001.1.region027 | 16 | *Streptomyces sp. CBMAI 2042* | Terpene singleton | NZ_RCOL01000001.1 | NCBI |
| NC_014228.1.region002 | 17 | *Xenorhabdus* | Beta-lactone | NC_014228.1 | NCBI |

| | | | | | |
|---|---|---|---|---|---|
| | | *nematophila ATCC 19061* | singleton | | |
| NZ_LXWF01000022.1.region002 | 18 | *Rothia nasimurium* | Beta-lactone singleton | GCF_002087015.1 | NCBI |
| NZ_CP045429.1.region006 | 19 | *Pseudoalteromonas rubra S4059* | T3PKS singleton | NZ_CP045429.1 | NCBI |
| NZ_RCOL01000001.1.region006 | 20 | *Streptomyces sp. CBMAI 2042* | T3PKS singleton | NZ_RCOL01000001.1 | NCBI |
| NZ_RCOL01000001.1.region016 | 21 | *Streptomyces sp. CBMAI 2042* | T1PKS singleton | NZ_RCOL01000001.1 | NCBI |
| NZ_RCOL01000001.1.region017 | 22 | *Streptomyces sp. CBMAI 2042* | lanthipeptide i singleton | NZ_RCOL01000001.1 | NCBI |
| NZ_LXWF01000041.1.region001 | 23 | *Rothia nasimurium* | NRP-metallophore singleton | GCF_002087015.1 | NCBI |

**Supplementary Table S5**. Average total runtime in seconds of BiG-SCAPE and BiG-SLiCE versions 1.1 and 2.0 runs, on random partitions of antiSMASH database of increasing size, as depicted in Figure 2.a. *The estimated runtime (see methods) of a single crashed BiG-SCAPE 1.1 is added to provide more information on how this version would continue to scale (Supplementary Fig. S8).

| Input dataset size (GenBank files) | BiG-SLiCE 1.1 | BiG-SLiCE 2.0 | BiG-SCAPE 1.1 | BiG-SCAPE 2.0 | Relative performance BiG-SCAPE (2.0 1.1) |
|---|---|---|---|---|---|
| 10 | 7.774 | 19.639 | 25.231 | 12.806 | 1.97 |
| 25 | 16.087 | 46.985 | 34.688 | 11.035 | 3.143 |
| 50 | 20.042 | 44.827 | 63.175 | 15.63 | 4.042 |
| 75 | 26.697 | 57.84 | 91.843 | 20.901 | 4.394 |
| 100 | 39.169 | 63.885 | 122.942 | 25.116 | 4.895 |
| 250 | 72.624 | 88.418 | 297.842 | 52.982 | 5.622 |
| 500 | 124.7 | 129.676 | 622.448 | 84.908 | 7.331 |
| 750 | 182.015 | 173.552 | 958.245 | 127.02 | 7.544 |
| 1000 | 211.759 | 209.108 | 1257.057 | 161.205 | 7.798 |
| 2500 | 514.83 | 478.01 | 3185.814 | 429.856 | 7.411 |
| 5000 | 1098.313 | 958.286 | 6710.19 | 1029.801 | 6.516 |
| 10000 | 2198.197 | 1845.769 | 14461.623 | 2947.241 | 4.907 |
| 25000 | 5387.304 | 4661.731 | 44968.949 | 19042.813 | 2.361 |
| 50000 | 17854.447 | 9146.052 | 153030* | 101310.839 | |
| 75000 | 18461.726 | 13017.656 | | 262954.296 | |

**Supplementary Table S6**. Average runtime of BiG-SCAPE 2.0, using the 10000 GenBank antiSMASH DB triplicate partitions, and alternative record types (*region*, *protocluster*, *protocore*), as well as alternative extension strategies (Table 1).

| **Variable to benchmark** | **Average time (seconds)** |
|---|---|
| Legacy extend strategy | 2640.283 |
| Greedy extend strategy | 2653.058 |
| Simple Match extend strategy | 2603.775 |
| Region record type | 2735.676 |
| Protocluster record type | 2985.594 |
| Protocore record type | 2866.841 |

**Supplementary Table S7.** Proportional and absolute runtimes of the major tasks (Input parsing, hmmscan, hmmalign, distance calculation and GCF calling) performed by BiG-SCAPE versions 1.1 and 2.0, at increasing input dataset sizes.

| | | Proportional Runtimes | | | | | Absolute Runtimes | | | | | |
|---|---|---|---|---|---|---|---|---|---|---|---|---|
| **Record count** | **version** | **Input Parsing** | **hmmscan** | **hmmalign** | **Distance calc** | **GCF calling** | **Input Parsing** | **hmmscan** | **hmmalign** | **Distance calc** | **GCF calling** | **Total runtime (s)** |
| 10 | v1.1 | 0,040 | 0,000 | 0,502 | 0,238 | 0,000 | 1,000 | 0 | 13 | 6 | 0 | 25 |
| 25 | v1.1 | 0,038 | 0,356 | 0,346 | 0,231 | 0,010 | 1,333 | 12 | 12 | 8 | 0 | 35 |
| 50 | v1.1 | 0,047 | 0,543 | 0,169 | 0,232 | 0,005 | 3,000 | 34 | 11 | 15 | 0 | 63 |
| 75 | v1.1 | 0,051 | 0,577 | 0,152 | 0,214 | 0,004 | 4,667 | 53 | 14 | 20 | 0 | 92 |
| 100 | v1.1 | 0,046 | 0,645 | 0,111 | 0,193 | 0,003 | 5,667 | 79 | 14 | 24 | 0 | 123 |
| 250 | v1.1 | 0,051 | 0,670 | 0,046 | 0,222 | 0,009 | 15,333 | 200 | 14 | 66 | 3 | 298 |
| 500 | v1.1 | 0,051 | 0,672 | 0,029 | 0,232 | 0,016 | 31,667 | 418 | 18 | 145 | 10 | 622 |
| 750 | v1.1 | 0,048 | 0,677 | 0,020 | 0,235 | 0,019 | 46,000 | 649 | 19 | 225 | 18 | 958 |
| 1000 | v1.1 | 0,045 | 0,668 | 0,018 | 0,244 | 0,024 | 56,333 | 840 | 22 | 307 | 31 | 1257 |
| 2500 | v1.1 | 0,045 | 0,666 | 0,013 | 0,232 | 0,044 | 144,667 | 2121 | 41 | 738 | 141 | 3186 |
| 5000 | v1.1 | 0,041 | 0,630 | 0,011 | 0,245 | 0,072 | 275,500 | 4230 | 75 | 1642 | 481 | 6710 |
| 10000 | v1.1 | 0,044 | 0,587 | 0,018 | 0,230 | 0,119 | 640,000 | 8493 | 267 | 3319 | 1727 | 14462 |
| 25000 | v1.1 | 0,035 | 0,461 | 0,031 | 0,234 | 0,239 | 1556,000 | 20753 | 1387 | 10532 | 10735 | 44969 |
| 10 | v2.0 | 0.069 | 0.277 | 0.135 | 0.037 | 0.483 | 0.879 | 3.541 | 1.728 | 0.471 | 6.187 | 12.806 |
| 25 | v2.0 | 0.113 | 0.529 | 0.171 | 0.038 | 0.149 | 1.246 | 5.838 | 1.889 | 0.417 | 1.645 | 11.035 |

| | | | | | | | | | | | | |
|---|---|---|---|---|---|---|---|---|---|---|---|---|
| 50 | v2.0 | 0.130 | 0.525 | 0.188 | 0.054 | 0.103 | 2.035 | 8.212 | 2.939 | 0.837 | 1.607 | 15.630 |
| 75 | v2.0 | 0.149 | 0.512 | 0.194 | 0.072 | 0.072 | 3.119 | 10.707 | 4.052 | 1.515 | 1.508 | 20.901 |
| 100 | v2.0 | 0.152 | 0.518 | 0.181 | 0.084 | 0.065 | 3.828 | 12.999 | 4.554 | 2.112 | 1.623 | 25.116 |
| 250 | v2.0 | 0.172 | 0.464 | 0.153 | 0.171 | 0.041 | 9.095 | 24.570 | 8.108 | 9.052 | 2.158 | 52.982 |
| 500 | v2.0 | 0.201 | 0.540 | 0.165 | 0.052 | 0.042 | 17.093 | 45.846 | 14.005 | 4.395 | 3.568 | 84.908 |
| 750 | v2.0 | 0.190 | 0.536 | 0.164 | 0.071 | 0.038 | 24.195 | 68.145 | 20.817 | 9.072 | 4.791 | 127.020 |
| 1000 | v2.0 | 0.182 | 0.526 | 0.161 | 0.091 | 0.040 | 29.419 | 84.727 | 26.024 | 14.664 | 6.371 | 161.205 |
| 2500 | v2.0 | 0.152 | 0.471 | 0.144 | 0.171 | 0.061 | 65.382 | 202.282 | 62.105 | 73.664 | 26.425 | 429.856 |
| 5000 | v2.0 | 0.120 | 0.380 | 0.121 | 0.286 | 0.092 | 123.938 | 391.785 | 124.894 | 294.297 | 94.888 | 1029.801 |
| 10000 | v2.0 | 0.076 | 0.260 | 0.083 | 0.446 | 0.134 | 223.407 | 767.246 | 245.862 | 1314.522 | 396.204 | 2947.241 |
| 25000 | v2.0 | 0.034 | 0.101 | 0.033 | 0.662 | 0.170 | 641.015 | 1932.833 | 619.103 | 12604.946 | 3244.916 | 19042.813 |
| 50000 | v2.0 | 0.012 | 0.041 | 0.012 | 0.790 | 0.144 | 1199.833 | 4198.515 | 1247.065 | 80057.173 | 14608.254 | 101310.839 |
| 75000 | v2.0 | 0.006 | 0.025 | 0.007 | 0.826 | 0.136 | 1507.050 | 6578.515 | 1868.496 | 217326.414 | 35673.821 | 262954.296 |
