## Supplementary Figures for "BiG-SCAPE 2.0 and BiG-SLiCE 2.0: scalable, accurate and interactive sequence clustering of metabolic gene clusters"

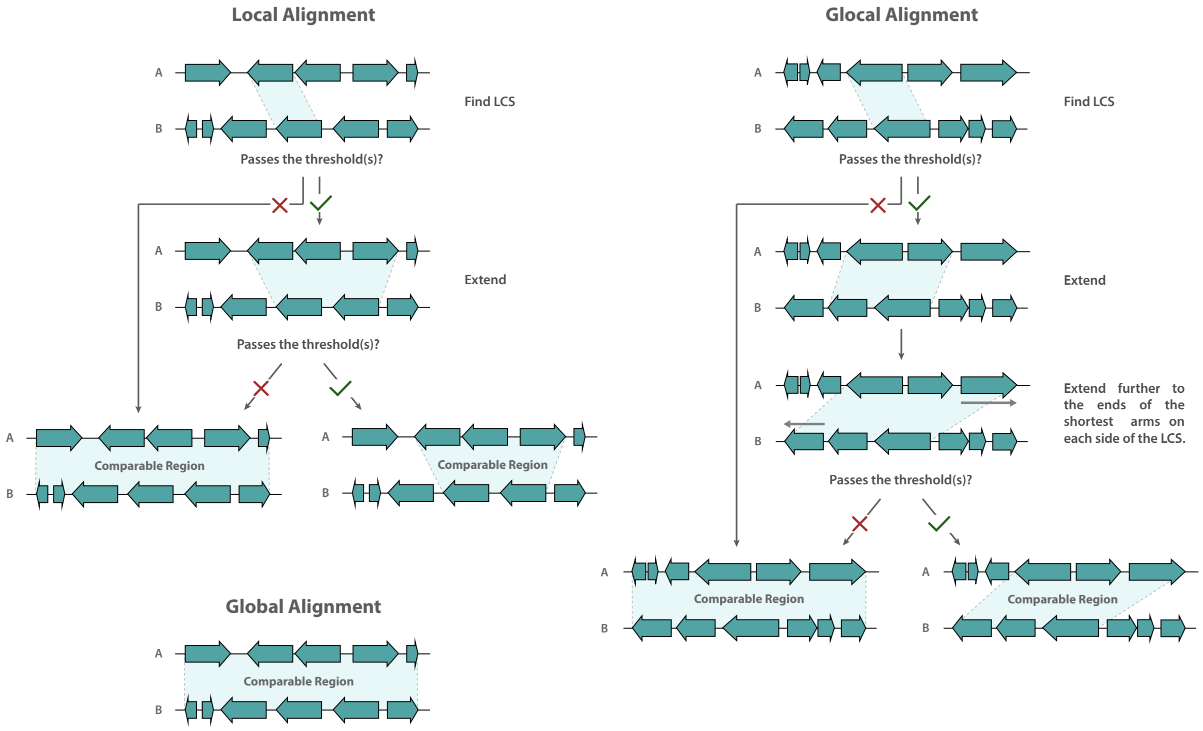


**Supplementary Figure S1.** Schematic overview of the behavior encoded in each of the alignment modes available in BiG-SCAPE 2.0. Local alignment consists of two stages: (i) finding a Longest Common Subsequence (LCS) and (ii) extending this LCS based on chosen extend strategy (Supplementary Fig. S3). Glocal alignment follows the same logic with an additional extension of the shortest arm on each side of the LCS. Both aforementioned modes rely on the checks described in Supplementary Figure S2. Global alignment does not rely on extension and directly compares the complete region.

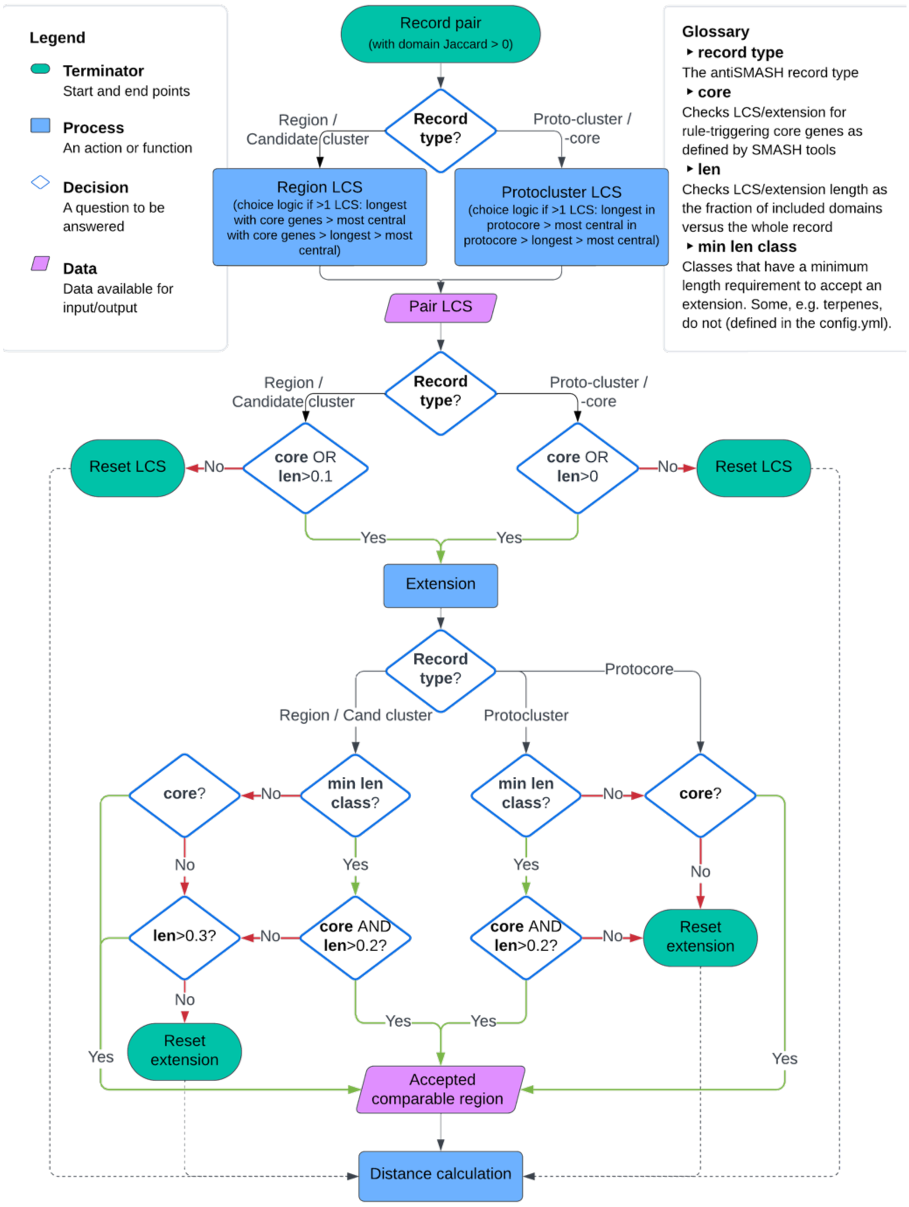


**Supplementary Figure S2.** Flowchart of processes and decisions carried out by BiG-SCAPE 2.0 in order to define the *comparable region* between a pair of BGC records. Decisions involving “core” refer to whether or not a relevant *comparable region* contains at least one domain annotated by antiSMASH, or related SMASH tools, as a biosynthetic/catabolic/rule-triggering domain. Similarly, “len” refers to the fraction of domains in the relevant region compared to the full BGC record. Throughout the process, biosynthetic content and relative length checks are performed to ensure that only a relevant LCS/*comparable region* is accepted. If these checks fail, the process is stopped and the comparable region of the BGC record pair defaults to the global alignment. Gene clusters not processed by SMASH tools will be handled as record type *region* and checks will only rely on relative length of the LCS/comparable region, since no “core” annotation will be present.

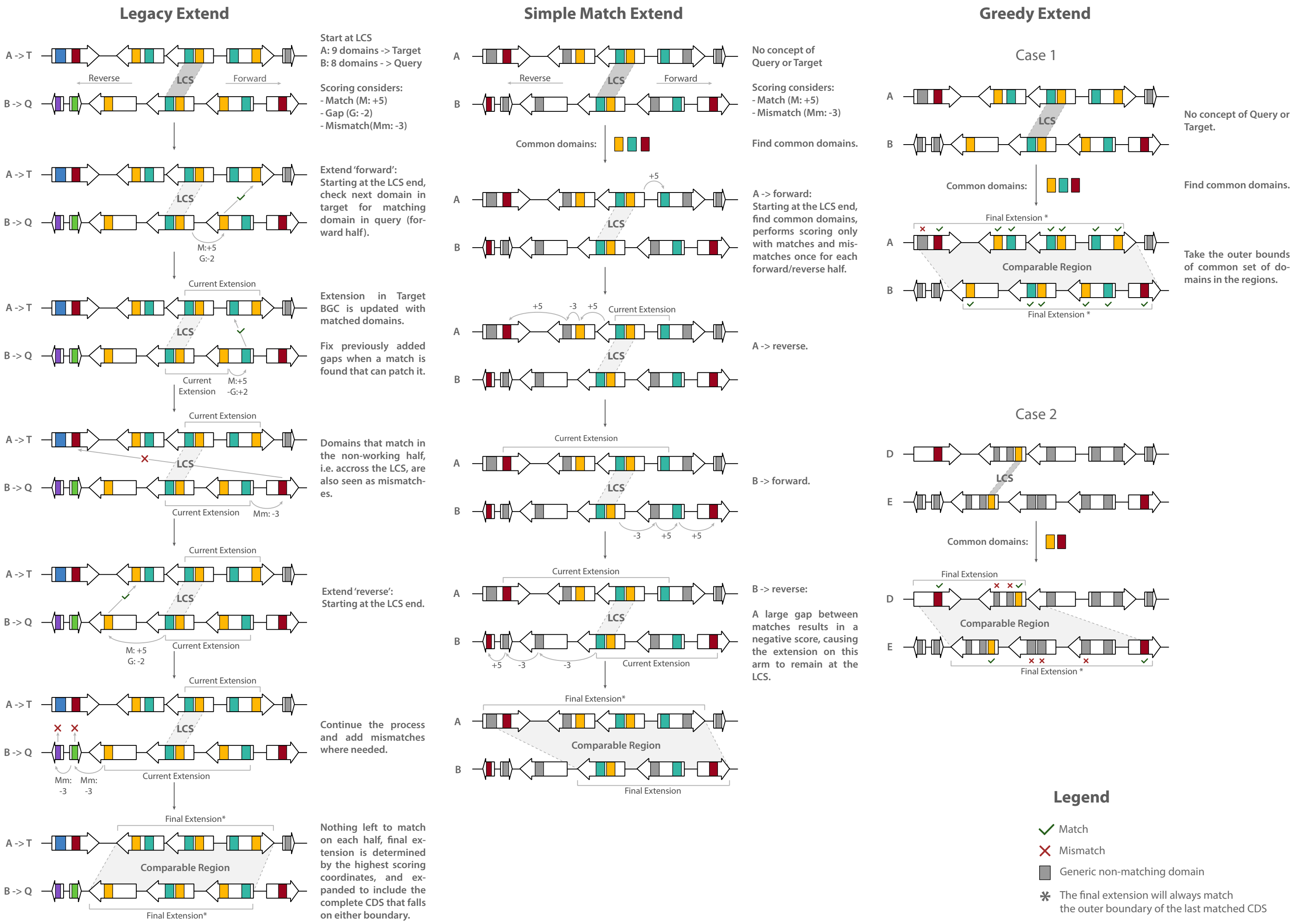


**Supplementary Figure S3.** Complete schematic representation of the extension strategies available in BiG-SCAPE 2.0. BiG-SCAPE 2.0 features three match/mismatch penalty algorithm extension strategies: Legacy Extend, Simple Match Extend and Greedy Extend. Legacy Extend follows the same principle used in BiG-SCAPE 1 (18); it defines a query (containing the fewest domains) and a target BGC record (if the two records have the same number of domains, query and target are randomly assigned). For each query domain, it subsequently searches for matching domains in the target. Legacy Extend is the strictest of the strategies, considering only matches on the same side (upstream or downstream) of the LCS, as well as gaps. Simple Match Extend has a higher tolerance for diverse regions, which performs the domain selection on all four arms of the BGC record pair, applying a match/mismatch scoring that does not consider domain positions/gaps. Greedy Extend is the simplest method, setting the coordinates of the comparable region at the first and last matching domains between the BGC record pair. In all cases, all domains of all CDSs at the edges of the comparable region are included.

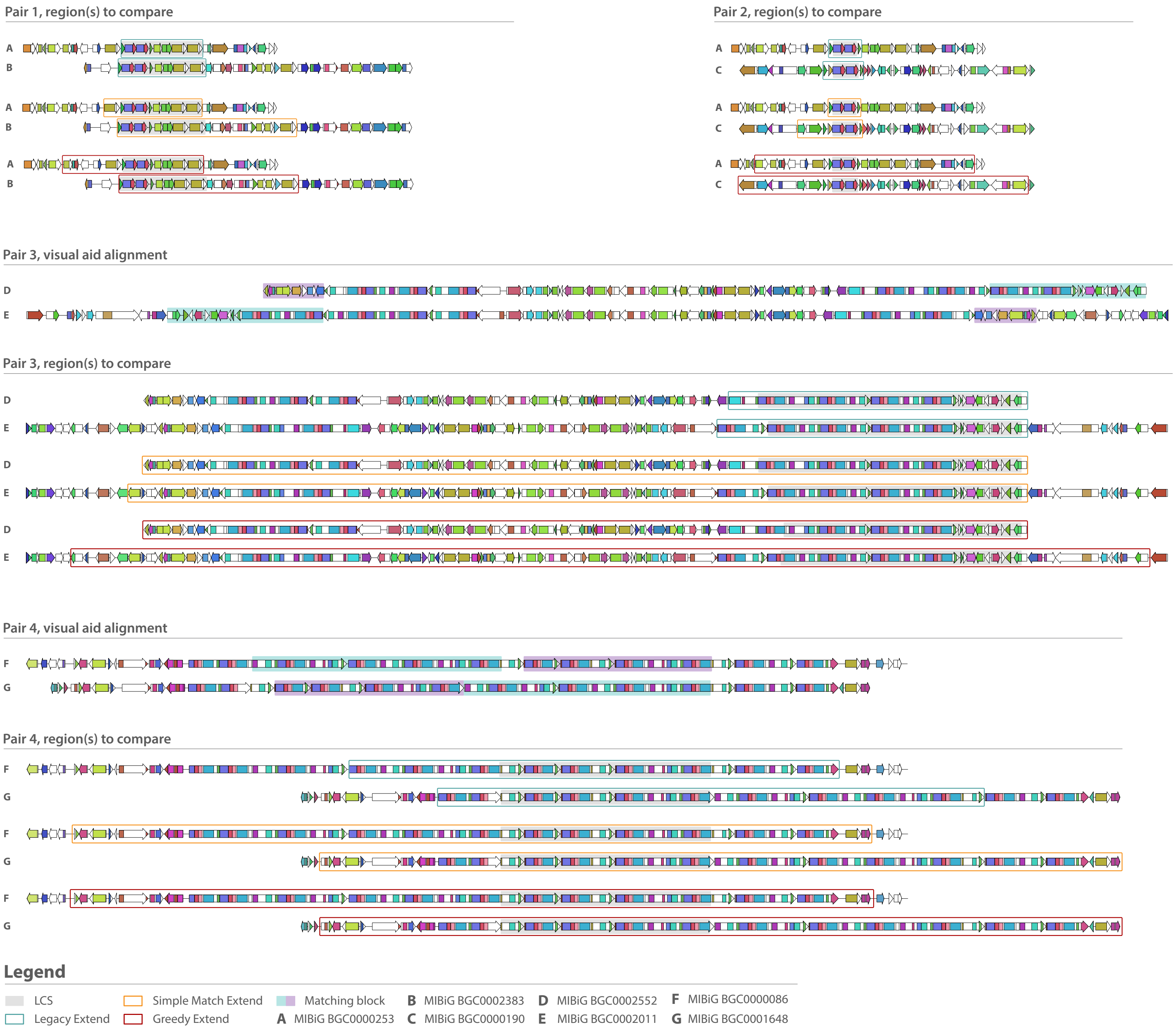


**Supplementary Figure S4**. Visual representation of computed comparable regions resulting from different extension strategies in four unique MIBiG BGC record pairs (Supplementary Table S1). Pair 1 and 2 represent straightforward examples where extension strategies can result in less or more similar comparable regions. More lenient extension strategies thus result in larger comparable regions. Pair 3 and 4 highlight extension behaviour when encountering translocated/recombinated regions within a BGC pair. Both simple match and greedy extend strategies rely less on domain position and are able to reach comparable regions more congruent with the visual aid alignment.

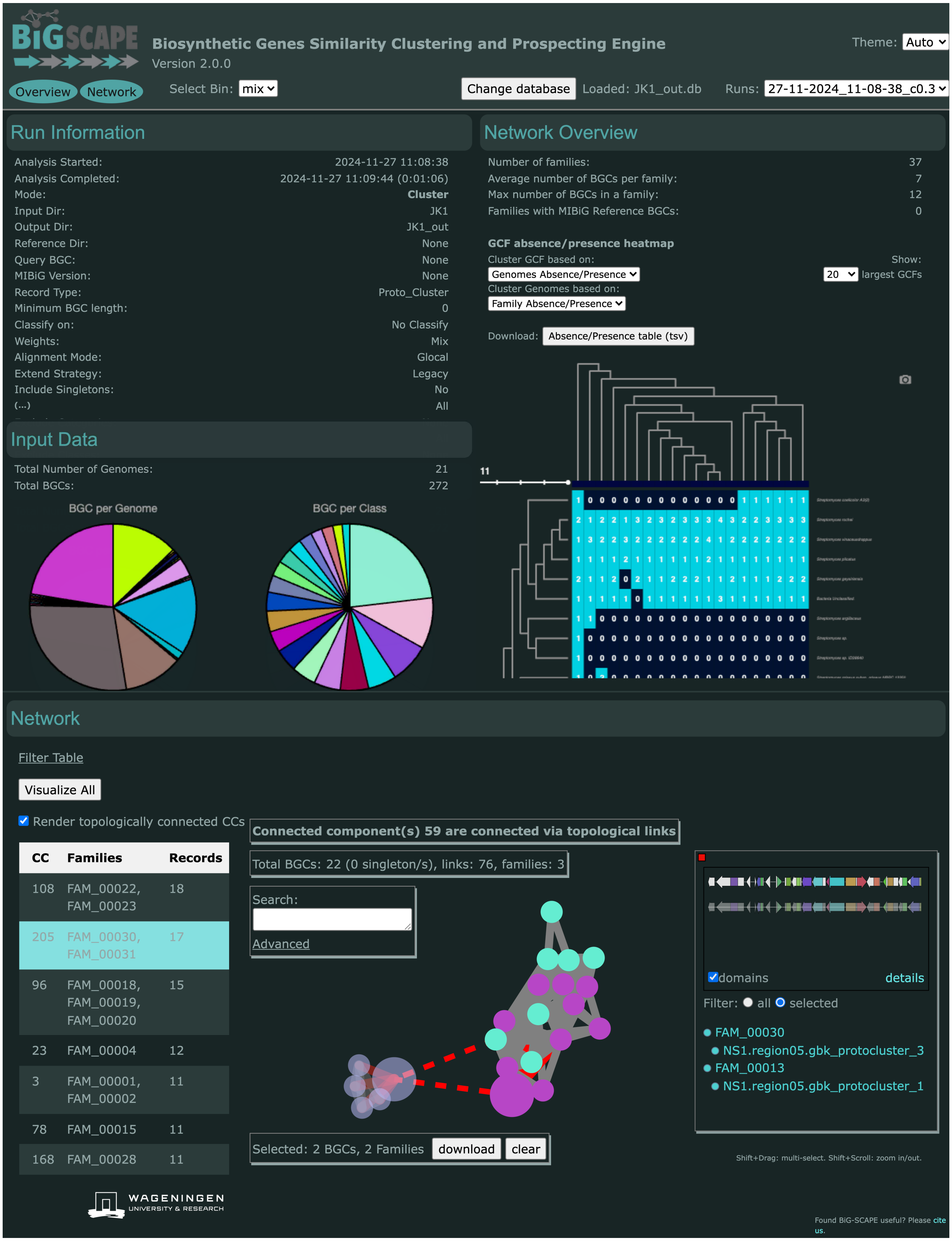


**Supplementary Figure S5.** Screenshot of BiG-SCAPE 2.0 user interface (UI) (run parameters: JK1 dataset, cluster mode, using the mix option, not classifying, record type *protocluster,* and otherwise all default parameters). BiG-SCAPE 2.0's interactive UI loads all required information from the output SQLite DB and populates a static index.html. The Network section has been significantly restructured and is organized on the basis of a connected component (CC) table. When a CC is selected its subnetwork will be loaded into view. Interacting with the loaded (sub)networks, e.g. selection of nodes and families, remains consistent with BiG-SCAPE version 1, as do the BGC detail and GCF tree panels (Supplementary Fig. S6). The CC table can be filtered by several BGC record specific elements, namely GenBank Description, name/path, pHMM/Pfam domain name and GCF identifier). For each CC's loaded network, the same filtering elements can be compounded in the advanced search box which handles logical operators, e.g. "[pHMM/Pfam domain name] AND [GCF identifier]". The filtered selections (at either CC or network level) can additionally be easily downloaded as .tsv files. To leverage the antiSMASH *region* concept, two types of links (edges) are now present, topological links and similarity links. Similarity links refer to the edges resulting from a calculated distance between two BGC records, and are represented as solid lines. Topological links refer to edges indicating that two BGC records were derived from the same antiSMASH *region*; and these are represented as dashed lines. Toggling the 'Render topologically connected CCs' button will load into the network view any CCs that are connected to the selected CC via topological links, which will be shown with semi transparent nodes and edges. In the node selection detail panel, as well as the GCF Tree view (Supplementary Fig. S6) the complete antiSMASH region is showcased, featuring a visual distinction between domains that belong to the relevant BGC record (in solid colors) and domains that belong to other BGC records within the region (in semi-transparent colors).

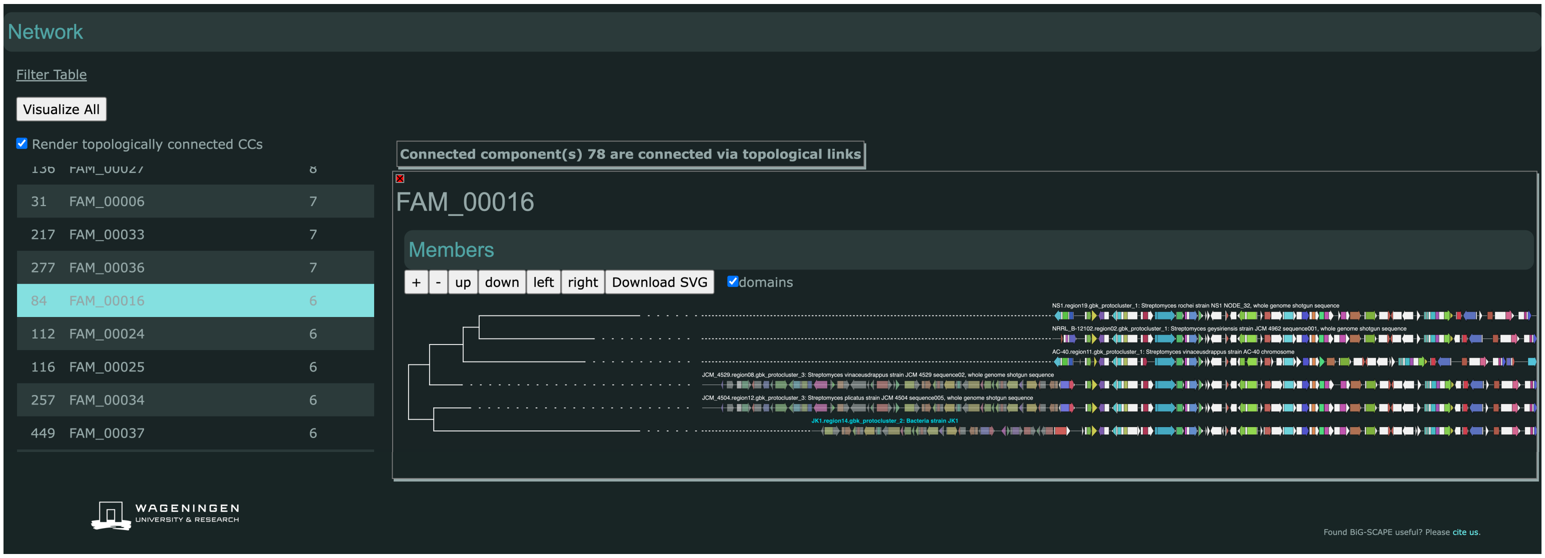


**Supplementary Figure S6**. Screenshot example of BiG-SCAPE 2.0 user interface GCF panel for FAM_00016 of run depicted in Supplementary Figure S5 (run parameters: JK1 dataset, run with cluster workflow, using the mix option, not classifying, record type *protocluster,* and otherwise with all default parameters). In this panel, the complete antiSMASH region is showcased, with domains that belong to the relevant BGC record displayed in solid colors, and domains that belong to other BGC records within the region displayed in semi-transparent colors.

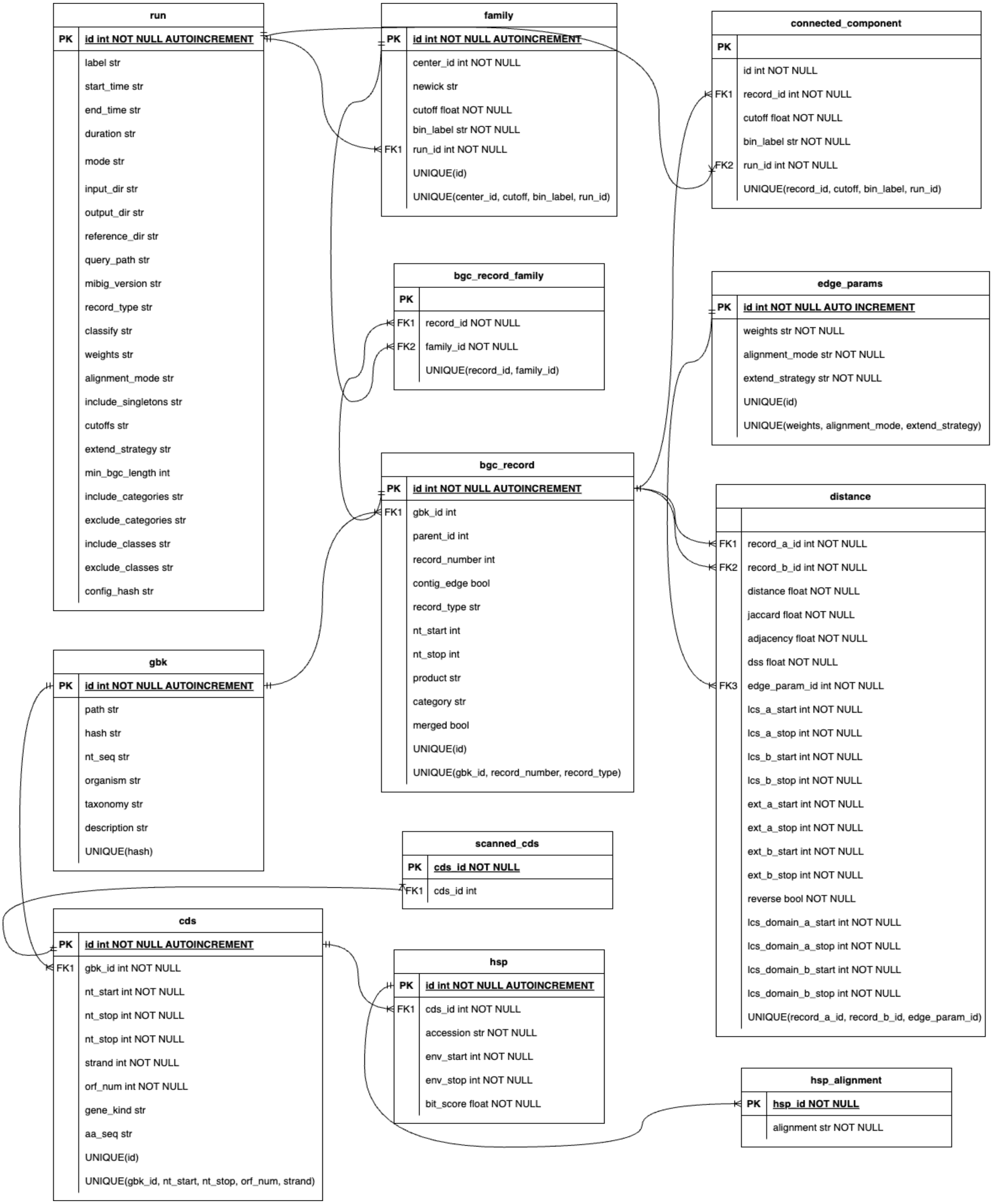


**Supplementary Figure S7**. BiG-SCAPE 2.0 Sqlite database schema.

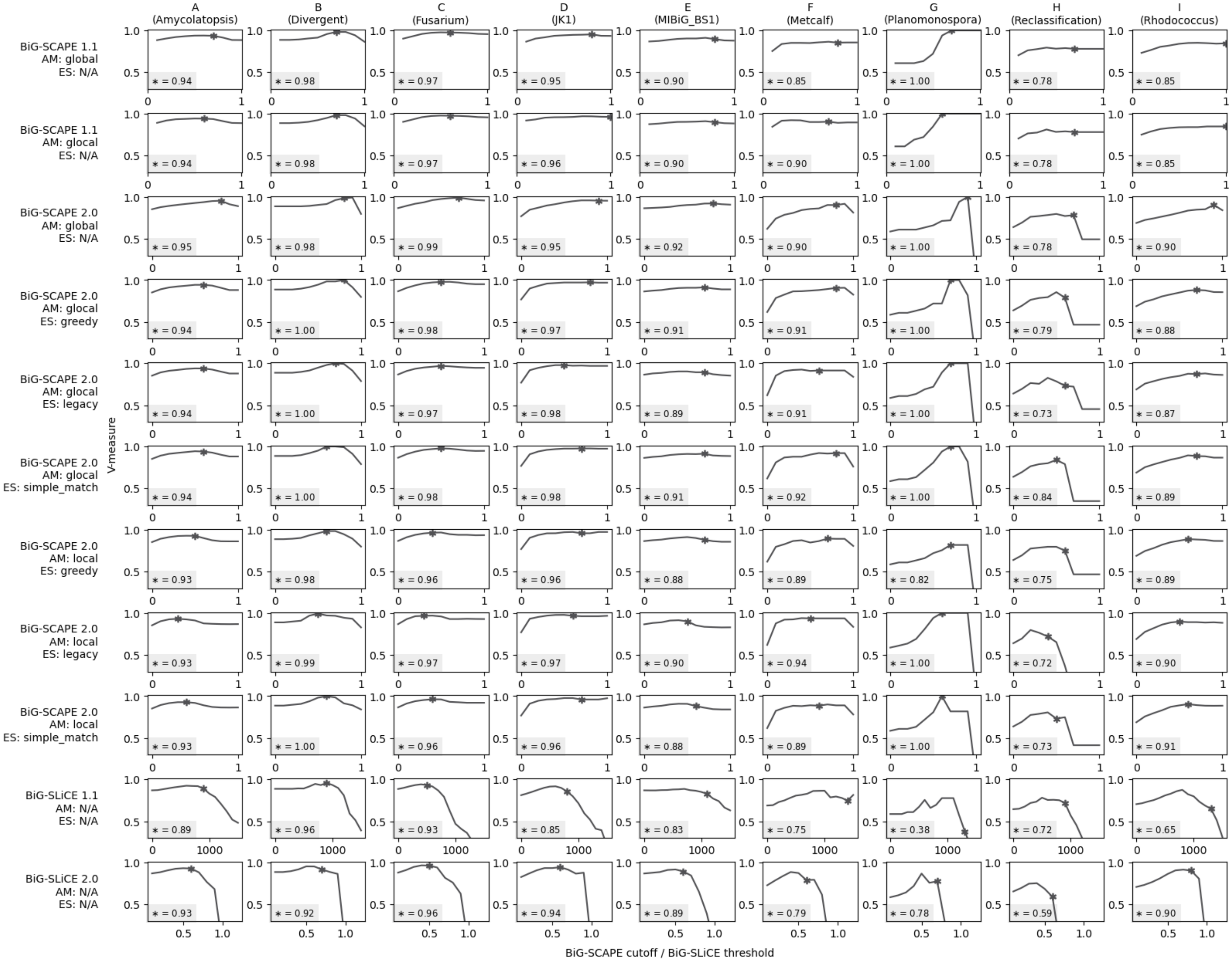


**Supplementary Figure S8**. A collection of graphs showing V-measure against BiG-SCAPE cutoff/BiG-SLiCE threshold. Columns in the grid refer to the input curated datasets A through I (Supplementary Table S6). Rows show the used tool version and, if applicable, BiG-SCAPE alignment mode (AM) and extension strategy (ES). In each graph, the optimal V-measure at the cutoff that results in a number of computed GCFs most similar to the number of curated GCFs is denoted by a star. The exact value is additionally shown in the bottom left of each graph.

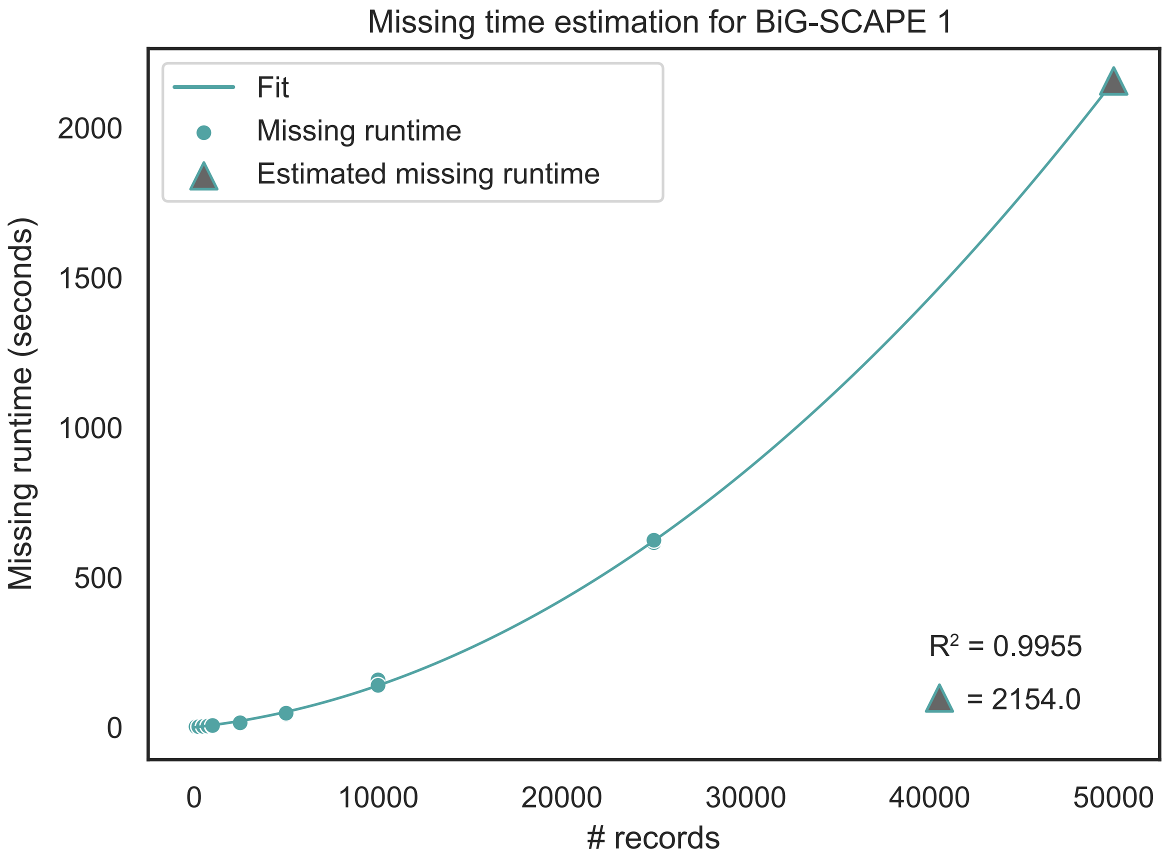


**Supplementary Figure S9.** Estimated ‘missing runtime’ of an incomplete BiG-SCAPE 1 run with an input of 50 000 BGC records, obtained by fitting a second order polynomial to the equivalent ‘missing runtimes’ for completed runs with inputs between 1 and 25 000 records. Missing runtimes are defined as the portion of the runtime between the creation of the last created file in the incomplete run, and the end of the total runtime.
